# The role of N-terminal acetylation on biomolecular condensation

**DOI:** 10.1101/2025.08.21.671499

**Authors:** Carolina G. Oliveira, Mayra T. S. Silva, Emanuel Kava, Tanushree Agarwal, Gea Cereghetti, Eduardo Festozo Vicente, Tuomas P. J. Knowles, Antonio J. Costa-Filho, Luis F. S. Mendes

**Affiliations:** Sao Carlos Institute of Physics, University of Sao Paulo, IFSC – USP, 13566-590, Sao Carlos, SP, Brazil; Molecular Biophysics Laboratory, Department of Physics, Faculty of Philosophy, Sciences, and Letters at Ribeirão Preto, University of São Paulo, Ribeirão Preto, SP, Brazil; Yusuf Hamied Department of Chemistry, Centre for Misfolding Diseases, University of Cambridge, Lensfield Road, Cambridge, CB2 1EW, UK; Cavendish Laboratory, Department of Physics, University of Cambridge, J J Thomson Ave, Cambridge, CB3 0HE, UK; Department of Biosystems Engineering, School of Science and Engineering, São Paulo State University (UNESP), Tupã, São Paulo, Brazil

**Keywords:** N-terminal Acetylation, Biomolecular Condensation, LLPS

## Abstract

N-terminal acetylation (Nt-acetylation) is one of the most prevalent co-translational modifications in eukaryotes, affecting nearly 80% of the human proteome. Despite its ubiquity, the potential impact of this phenomenon on biomolecular condensation has been largely overlooked. Here, we uncover how this chemically subtle modification can exert broad and multifaceted control over phase behaviour, using Grh1, a Golgi-associated protein involved in stress-induced secretion in yeast, as a model system. We show that Nt-acetylation increases the saturation concentration for condensation, reduces droplet size and number, dampens pH sensitivity, weakens electrostatic contributions, and suppresses water dipolar relaxation within condensates, indicating reduced internal hydration and environmental responsiveness. These effects are accompanied by acetylation-dependent dimerisation and local structural changes, including a concentration-dependent gain in α-helicity. Remarkably, co-condensation assays reveal that acetylated and non-acetylated forms of the same protein are only partially miscible, giving rise to core-shell architectures driven by differences in interfacial tension. Together, our findings highlight Nt-acetylation as a potent, generalizable regulator of condensate material properties, linking primary sequence chemistry to mesoscale organisation. Given its evolutionary conservation and prevalence across eukaryotic proteomes, Nt-acetylation may represent a widespread mechanism for modulating protein condensation in health and disease.

## Introduction

Biomolecular condensation is a widespread and essential phenomenon in cell biology, enabling the formation of membraneless compartments that organise biochemical reactions in space and time ^1,2^. This process, known as phase separation coupled to percolation (PSCP), relies on weak, multivalent interactions among macromolecules, especially proteins containing intrinsically disordered regions (IDRs) ^2–5^. Biomolecular condensation drives the formation of dynamic, reversible, non-stoichiometric droplets involved in key cellular processes such as transcriptional regulation, RNA metabolism, protein quality control, the organisation of the cytoskeleton, and stress response ^6–8^.

Several paradigmatic examples highlight the biological importance of protein condensation. In the nucleus, biomolecular condensation underlies the organisation of the nucleolus and heterochromatin domains, with proteins such as NPM1 and HP1α forming distinct coexisting phases to regulate rRNA processing and gene silencing, respectively ^9–13^. In the cytoplasm, biomolecular condensation governs the assembly of stress granules and P-bodies, transient compartments that sequester untranslated mRNAs and translation factors during environmental stress ^14,15^. Even viral replication and innate immune responses have been shown to depend on condensate formation, as exemplified by the mitochondrial antiviral protein MAVS and the cGAS–STING pathway ^16,17^. These functions are possible because condensates concentrate specific biomolecules, enhance reaction kinetics, and enable spatial segregation of incompatible processes.

Although the propensity for biomolecular condensation is encoded in sequence features such as low complexity, charge patterning, and domain valency, there is growing recognition that post-translational modifications (PTMs) play a central role in regulating condensate formation, dissolution, and material properties ^18–20^. PTMs such as phosphorylation, methylation, acetylation, and ubiquitination can modulate phase separation by altering electrostatics, hydrophobicity, structural flexibility, and interaction networks ^18,21,22^. Several examples illustrate this regulatory power. Phosphorylation of FUS prevents its pathological transition into solid aggregates, preserving a liquid-like condensate state ^22,23^. Arginine methylation of FUS and DDX4 suppresses condensation by interfering with cation-π interactions critical for condensate formation ^24,25^. UBQLN2-mediated phase separation selectively protects K48-linked polyubiquitinated proteins from deubiquitination, highlighting a mechanism for regulating protein stability ^26^. Lysine acetylation of DDX3X modulates its phase separation properties, suggesting a regulatory role for this post-translational modification in RNA granule assembly and neurodevelopmental function ^27,28^.

Among these better-studied modifications, N-terminal acetylation (Nt-acetylation) remains surprisingly underappreciated despite its ubiquity. Nt-acetylation is one of the most common PTMs in eukaryotes, present in approximately 80% of the human proteome ^29,30^. Catalysed by N-terminal acetyltransferases (NATs), this irreversible reaction transfers an acetyl group from acetyl-CoA to the free α-amino group of the N-terminal residue, typically occurring co-translationally as the nascent polypeptide emerges from the ribosome ^30^. This modification neutralises the positive charge of the N-terminal amine and introduces a chemically distinct cap, influencing the polarity, rigidity, and interaction potential of the protein’s N-terminus, although modestly ^30^.

Functionally, Nt-acetylation has been linked to numerous cellular roles, including stabilising α-helical structures via the helix dipole effect, regulating protein half-life and degradation through N-degron pathways, modulating membrane association, and affecting protein-protein interactions ^30^. Importantly, it often operates in context-dependent ways, with the same modification producing opposing effects in different protein environments ^30^. Yet, the implications of this PTM in biomolecular condensation remain largely unexplored, particularly in *in vitro* studies that use recombinant proteins lacking native acetylation patterns, which may misrepresent physiological biomolecular condensation behaviour.

In this study, we address this gap by investigating the role of Nt-acetylation in modulating the phase behaviour of Grh1 (Golgi Reassembly and Stacking Protein Homologue 1), a yeast GRASP family member involved in Golgi organisation and unconventional protein secretion ^31,32^. Grh1 features a conserved N-terminal with PDZ domains and a disordered C-terminal SPR (serine–proline-rich) domain, making it prone to biomolecular condensation ^33–35^. Here, we produced acetylated and non-acetylated forms of Grh1 through co-expression with the yeast NatC complex and compared their behaviour using biophysical and structural assays. Our findings reveal a dramatic impact of N-terminal acetylation on condensate formation, morphology, rheology, and saturation threshold. These results position Nt-acetylation as a powerful yet frequently neglected modulator of phase separation. They also underscore the necessity of accounting for PTMs, particularly irreversible and co-translational modifications, such as N-terminal acetylation, in future biomolecular condensation studies for accurate experimental design and biological interpretation.

## Methods

### Protein expression and purification

For heterologous expression in bacteria, plasmid vectors carrying the genes of interest were introduced into *E. coli* Rosetta competent cells. Grh1 was cloned into the pET22b(+) vector and purified following an adapted protocol from previous studies ^33,36^. Briefly, transformed cells were grown in LB medium at 37 °C with shaking at 200 rpm until an optical density (OD₆₀₀) of 0.8 was reached. Protein expression was induced by adding 500 µM isopropyl β-D-1-thiogalactopyranoside (IPTG), followed by incubation for 18 hours at 18 °C and 220 rpm. Cells were harvested by centrifugation at 8,000×g for 10 minutes and resuspended in 20 mL of buffer A (10 mM HEPES/NaOH, 300 mM NaCl, 5 mM 2-mercaptoethanol, 10% glycerol, pH 8.0) per litre of culture. Cell lysis was performed by sonication on ice using a Branson 450 Digital Sonifier® (72 pulses of 5 s at 20% amplitude with 10 s intervals). Lysates were clarified by centrifugation at 10,000×g for 20 minutes, and the supernatant was applied to a Ni²⁺-NTA affinity column pre-equilibrated with buffer A. After washing with 20 mL of buffer A and sequential imidazole steps (20 mM and 40 mM), the bound protein was eluted with 500 mM imidazole in buffer A. The eluate was concentrated using an Amicon Ultra-15 centrifugal filter unit (10 kDa NMWL, Merck Millipore) and further purified by size-exclusion chromatography using a Superdex 200 10/300 GL column coupled to an ÄKTA Purifier system (GE Healthcare Life Sciences). Acetylated Grh1 (AcGrh1) was obtained by co-expression of the Grh1-pET22b construct with the *Saccharomyces cerevisiae* NatC complex (full-length Naa30, Naa35, and Naa38) cloned into the pRSFDuet-1 vector (kindly provided by Prof. Oliver Daumke, MDC Berlin) ^37^. The NatC complex lacks an affinity tag and is completely removed during Ni-NTA purification. The same purification protocol was applied to isolate AcGrh1. For fluorescence-based analyses, purified proteins were labelled with Alexa Fluor 488 or Alexa Fluor 555 dyes (ThermoFisher). Dyes were prepared as 10 mM stock solutions in DMSO and added to protein samples at a twofold molar excess. Labelling reactions were conducted at room temperature (25 °C) for 1 hour under gentle agitation. Excess unbound dye was removed by size-exclusion chromatography, yielding purified, dye-labelled protein.

### Matrix-assisted laser desorption/ionisation mass spectrometry (MALDI-MS)

For mass spectrometry analysis, buffer exchange was performed under nondenaturing conditions to remove nonvolatile salts from Grh1 and AcGrh1 protein samples. The exchange to 10 mM ammonium acetate buffer (pH 7.4) was carried out using a HiTrap Desalting chromatography column (GE Healthcare), coupled to an ÄKTA Purifier system (GE Healthcare), at a flow rate of 1 mL/min. Following buffer exchange, samples were subjected to enzymatic digestion using Glutamyl endoproteinase from *Staphylococcus aureus* V8 (Glu-C) (Sigma-Aldrich). For each digestion reaction, 29 µL of ultrapure water and 50 µL of 100 mM ammonium acetate (pH 4.5) were added to 10 µL (1– 50 ng protein) of each protein sample. Dithiothreitol (DTT, 500 mM) was then added to a final concentration of 10 mM. Samples were gently mixed and incubated at 60 °C for 45 minutes. Subsequently, iodoacetamide (IAA, 500 mM) was added to a final concentration of 30 mM. The reaction mixtures were briefly centrifuged and incubated in the dark at room temperature for 30 minutes. To quench the alkylation reaction, DTT was added again to a final concentration of 20 mM. 0.5-2.5 µL Glu-C was then added at an enzyme-to-protein ratio of 1:20 (w/w). Samples were briefly mixed and incubated at 37 °C for 24 hours. After digestion, samples were flash-frozen in liquid nitrogen (N_2_) and stored at –80 °C until analysis. Before mass spectrometry, peptide cleanup was performed using C18 ZipTip pipette tips protocol (Millipore) to remove salts and concentrate the peptides. Mass spectra were acquired using a MALDI TOF/TOF mass spectrometer (UltrafleXtreme model, Bruker Daltonics GmbH & Co.).

### Circular Dichroism

Circular dichroism (CD) measurements were performed using a J-815 spectropolarimeter (Jasco), equipped with a Peltier temperature control system and a quartz cuvette with a 1 mm pathlength. Far-UV CD spectra were recorded in the 195–270 nm range, using a scanning speed of 50 nm min⁻¹, a spectral bandwidth of 1 nm, and a digital integration time of 2 s. Protein samples were diluted in 20 mM sodium phosphate buffer (pH 7.4) to a final concentration of 0.1 mg/mL, representing at least a 20-fold dilution from the stock solutions. The protein concentration was adjusted to 4 mg/mL for near-UV CD measurements. Spectra were collected in the 240–350 nm range under the same scanning speed (50 nm min⁻¹), with a spectral bandwidth of 2 nm and a digital integration time of 2 s. All experiments were performed in triplicate to ensure reproducibility. To ensure measurement reliability, a high-tension (HT) voltage threshold of 600 V was set as the cut-off criterion for both far- and near-ultraviolet (UV) spectral data. The data were processed and analysed using OriginPro^®^ 2022.

### Differential Scanning Calorimetry

Differential scanning calorimetry (DSC) measurements were performed using a VP-DSC MicroCal calorimeter (MicroCal, Northampton, MA, USA). All experiments were conducted at a constant heating rate of 1 °C/min over a temperature range of 20-70°C under a controlled pressure of 1.6 atm. Instrumental baselines were recorded using buffer-only runs before sample analysis to account for the thermal history of the system. Raw thermograms were baseline-corrected by subtracting the corresponding buffer trace and subsequently normalised by protein concentration (4 mg/mL). Thermal baselines (cubic, progressive, or manual) were reconstructed and subtracted from the normalised traces using MicroCal Origin software. Each experiment was independently repeated at least twice to ensure data reproducibility. Different denaturation models were fitted to the experimental DSC data using CalFitter 2.0 ^38^.

### Dynamic Light Scattering

The size distribution of protein condensates was analysed by Dynamic Light Scattering (DLS) using a Zetasizer Nano ZS (Malvern Panalytical, UK). Measurements were performed at 25°C in 20 mM sodium phosphate buffer (pH 5.4) supplemented with 10% (w/v) PEG-8000, which matched the conditions used for in vitro condensate formation. Protein solutions were prepared at final concentrations of 0.5 g/L and 1 g/L for both Grh1 and AcGrh1 for the phase-separated samples. For non-phase-separated conditions, samples were analysed in a working buffer alone at protein concentrations of 2 mg/mL and 10 mg/mL. Data acquisition was performed in backscatter mode at a detection angle of 173°, with 10 consecutive measurements of 10 seconds each. Data were analysed using Zetasizer Software v7.13. The cumulants method was applied to determine the mean hydrodynamic diameter, and size distributions were extracted based on the number to minimise the bias introduced by larger particles in the scattering intensity.

### Molecular Dynamics Simulation

The structure of the Grh1 N-terminal region (residues 1–28) was predicted using AlphaFold 3.0. Molecular dynamics simulations in explicit water were performed using the WebGro server (https://simlab.uams.edu/ProteinInWater/index.html; accessed in 2025). Simulations employed the GROMACS force field 96 43a1, the TIP4P water model, and a triclinic box with 150 mM NaCl. The systems were equilibrated and simulated at 303 K and 1 bar for 50 ns. Three independent simulations were carried out for each construct—acetylated and non-acetylated—and the results presented correspond to the combined trajectories. In all cases, 50 ns was sufficient for the system to converge to a fully unfolded conformation. Simulation outputs were analysed using VMD and CCP4MG.

### Size-exclusion chromatography coupled to multi-angle light scattering (SEC-MALS)

Size-exclusion chromatography coupled with multi-angle light scattering (SEC-MALS) was performed using a miniDAWN TREOS detector (Wyatt Technology, CA) equipped with three detection angles (43.6°, 90°, and 136.4°) and a 659 nm laser. A Wyatt QELS module was used in-line for dynamic light scattering (DLS) measurements to determine the hydrodynamic radius (R_H_), along with an Optilab® rEX refractive index detector (Wyatt Technology). All measurements were carried out using a Superdex 200 HR 10/300 analytical SEC column (GE Healthcare). Bovine serum albumin (BSA; Sigma-Aldrich) was used as a reference standard. Protein samples (∼2 mg/mL) were eluted in 20 mM Tris-HCl, 500 mM NaCl, pH 8.0, at a constant flow rate of 0.5 mL/min. Immediately before data acquisition, the samples were centrifuged at 10,000 × g for 10 minutes at 4°C to remove aggregates or debris. Data acquisition and analysis were performed using ASTRA 7 software (Wyatt Technology) with the following parameters: refractive index of 1.331, solvent viscosity of 0.890 cP, and a refractive index increment (dn/dc) of 0.185 mL/g.

### Turbidity assays

Turbidity measurements were performed using a Multiskan Go spectrophotometer (Thermo Fisher) operated with SkanIT RE 6.1.1 software, employing 96-well tissue culture plates (Kasvi). The samples consisted of unlabeled protein and the corresponding solution, composed of Na₂HPO₄/NaH₂PO₄ at the specified pH and supplemented with 10% (w/v) PEG 8000. All experiments were conducted in triplicate, and each sample was measured five times at an absorbance wavelength of 400 nm.

### Microscopy and Image Analysis

Biomolecular condensates formed by Grh1 and AcGrh1 were visualised using wide-field (DIC and fluorescence) and confocal microscopies. Differential interference contrast (DIC) and epifluorescence images were acquired on an Olympus IX71 inverted microscope equipped with a PicoQuant MT 200 module (PicoQuant®, Germany) and a 60× water-immersion objective lens (numerical aperture = 1.2). Fluorescence excitation was controlled using the following mirror units: U-MNU2 BP360/370 (UV), U-MWB2 BP460/490 (blue), and U-MWIG3 BP530–550 (green), with long-pass filters HQ405LP, HQ460LP, BLP01-488R, HQ550LP, and HQ690/70m. Protein samples were prepared in 20 mM sodium phosphate buffer (pH adjusted as needed), supplemented with 2–20% (w/v) PEG 8000 (ThermoFisher), and incubated at room temperature for 20 minutes before imaging. A 5 μL droplet of the solution was placed on a 20 × 20 mm coverslip (Deckgläser) for microscopy. Data acquisition was performed using SymPhoTime 64 (PicoQuant), and image analysis was conducted in ImageJ. Confocal fluorescence microscopy was performed using a Leica Stellaris 8 system equipped with a scanning laser and a HyD (Hybrid Detector) module. Samples of Grh1 and AcGrh1 labelled with Alexa Fluor 488 (green) and Alexa Fluor 546 (red), respectively, were prepared in 20 mM phosphate buffer at pH 5.4 containing 10% PEG 8000, with each protein at a final concentration of 1 mg/mL. To reduce surface interactions, samples were mounted on silanized 20 × 20 mm coverslips (Deckgläser) treated with mPEG5k. Image acquisition was performed using a 63× water-immersion objective (numerical aperture, NA 1.2). Excitation wavelengths were set to 488 nm and 561 nm, with emission collected in the 500–550 nm and 570–620 nm spectral windows, respectively. Three-dimensional reconstructions were obtained by Z-stack scanning at 0.3 μm intervals using a 512 × 512-pixel resolution. Spectral compensation and background subtraction were performed automatically with Leica LAS X software. The LUT (Look-Up Tables) plugin in Fiji/ImageJ was used for fluorescence intensity and colocalisation analyses.

### Hyperspectral imaging

Hyperspectral images were acquired using a confocal Leica Stellaris 5 microscope equipped with a HC PL APO CS2 63x/1.40 oil immersion objective (Leica Microsystems). A laser wavelength of 405 nm was used for ACDAN excitation. The ACDAN dissolved in DMSO was directly added to the unlabeled protein solution (50 µM protein concentration in 20 mM sodium phosphate, 7% PEG 8K, pH 5.4 or 7.4) before the experiment, to a final ACDAN concentration of 5 µM. Image acquisition was performed with a frame size of 512 × 512 pixels and a pixel size of 150 nm. The xyλ configuration of the microscope was used, sequentially measuring in 32 channels in the range from 415 to 725 nm. The images were processed using the SimFCS software, developed at the Laboratory of Fluorescence Dynamics, which is available on the webpage (https://www.lfd.uci.edu/globals/ accessed in 2025). The spectral phasor analysis utilises a Fourier transformation applied to the fluorescence emission spectra, where we calculate two parameters, G and S (as shown in the equations below), that represent the real and imaginary components of the transformation.

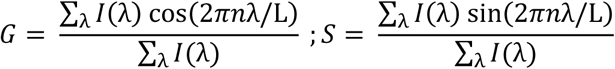

In the equations above, *I(λ)* is the intensity in each step of spectral measurement, *L* is the spectrum range (λ_max_ - λ_min_), and *n* is the harmonic number (=1 in this case). The spectral phasor plot consists of the two parameters G and S plotted in a four-quadrant polar coordinate system. In this case, the wavelength range is captured at each pixel of the hyperspectral image.

### Saturating concentration

Saturating concentrations were obtained by preparing 5× dilutions of the protein stock solution for all final concentrations used in the assay. This approach was designed to minimise potential interference from NaCl present in buffer A of the stock solution during condensate formation. All samples were prepared in 20 mM Na₂HPO₄/NaH₂PO₄ buffer supplemented with 10% (w/v) polyethylene glycol (PEG 8000). After sample preparation, the mixtures were centrifuged at 15,000 rpm for 15 minutes to separate the dense and diluted phases. The supernatant, corresponding to the diluted phase, was collected and analysed using a NanoDrop 2000/2000c spectrophotometer (ThermoFisher) by measuring absorbance at 280 nm to quantify the protein concentration (ε₂₈₀ = 32,890 M⁻¹.cm⁻¹). All measurements were performed in triplicate.

### Microfluidic device preparation

The microfluidic devices used at the PhaseScan platform were prepared using standard lithography processes ^39^. The devices were designed using AutoCAD software (Autodesk) and then printed on a photomask (Micro Lithography). A 45 µm layer of SU8-3050 photoresist (Microchem) was deposited on a silicon wafer using a spin coater with the following parameters: 10 s at 500 rpm, followed by 45 s at 2700 rpm with a 300 rpm/s acceleration. The coated wafers were heated to 95 °C for 20 minutes. The photomask was aligned on top of the wafer, and the pattern was transferred through a 1.5-minute UV exposure followed by a bake at 95 °C for 20 minutes. Excess unreacted SU8-3050 was removed using propylene glycol methyl ether acetate (PGMEA). The excess Polydimethylsiloxane (PDMS) was prepared at a 10:1 base-to-curing agent ratio, poured over the mask, and cured at 65 °C for 1.5 hours. After curing, the PDMS was cleaned, and holes for the inlet/outlet were punched. The PDMS layer was then plasma-bonded to glass slides for 30 s at 60% power using a plasma oven (Diener Electronics). The interior surfaces were treated with 1% trichloro(1H,1H,2H,2H-perfluorooctyl)silane (Sigma) in HFE-7500 (Fluorochem), then heated at 95 °C for 10 minutes.

### Phase diagrams acquisition

Phase Scan, a high-throughput microfluidic technique, was used to construct the phase diagrams ^39^. Ghr1 or AcGrh1 solutions containing 5 mol% of AlexaFluor 647-labelled protein were mixed at different mass ratios in buffer and encapsulated in water-in-oil droplets using the microfluidic device. For the protein vs. PEG concentration experiments, 10 mM Hepes, 300 mM NaCl, pH 7.4, was used as the buffer. For the pH scan experiments, the acidic and basic buffers used were 200 mM sodium phosphate and 200 mM succinic acid at pH levels of 3.5 and 9. The flow profile mixed the solutions, with 50% of the droplets containing a protein solution and the remaining 50% consisting of a buffer mixture. The expected pH values inside each droplet were estimated by a pH calibration curve obtained by mixing the acidic and basic buffers in different ratios ^40^. Fluorescence images of droplets within the microfluidic device’s imaging chamber were acquired using an epifluorescence microscope (Cairn Research) equipped with a 10× objective (Nikon CFI Plan Fluor 10×, NA 0.3). An automated image analysis script was used to detect condensates and quantify protein concentrations in individual droplets. The resulting data were visualised as scatter plots, displaying average values across experimental conditions.

## Results and Discussion

### Construction of the Nt-acetylated version of Grh1 (AcGrh1)

Grh1 is the yeast orthologue of mammalian GRASP65, and its N-terminal sequence begins with the motif MFRI, which makes it a substrate for the NatC N-terminal acetyltransferase complex. Acetylation is a post-translational modification essential for the membrane association of Grh1 ^31^, serving as a functional replacement for N-myristoylation, which is typically observed in GRASPs from metazoans ^41–43^. We adopted a previously established co-expression system to reconstitute this post-translational modification in *Escherichia coli*. A modified pRSFDuet-1 vector harbouring untagged full-length *S. cerevisiae* Naa30, Naa35, and Naa38, kindly provided by Prof. Oliver Daumke (Max Delbrück Center for Molecular Medicine, MDC), was used as described by Varga et al. ^37^. This vector was co-transformed with a pET22b plasmid encoding the C-terminally His-tagged Grh1 with a free N-terminus, allowing for co-translational acetylation during bacterial expression. Interestingly, differences between the two expression systems were already evident at the purification stage, with the acetylated version yielding significantly higher amounts of soluble protein than its non-acetylated counterpart. Successful Grh1 acetylation was confirmed by MALDI-MS, which indicated near-complete N-terminal modification using this co-expression protocol (Figure 1A).

**Figure 1:**
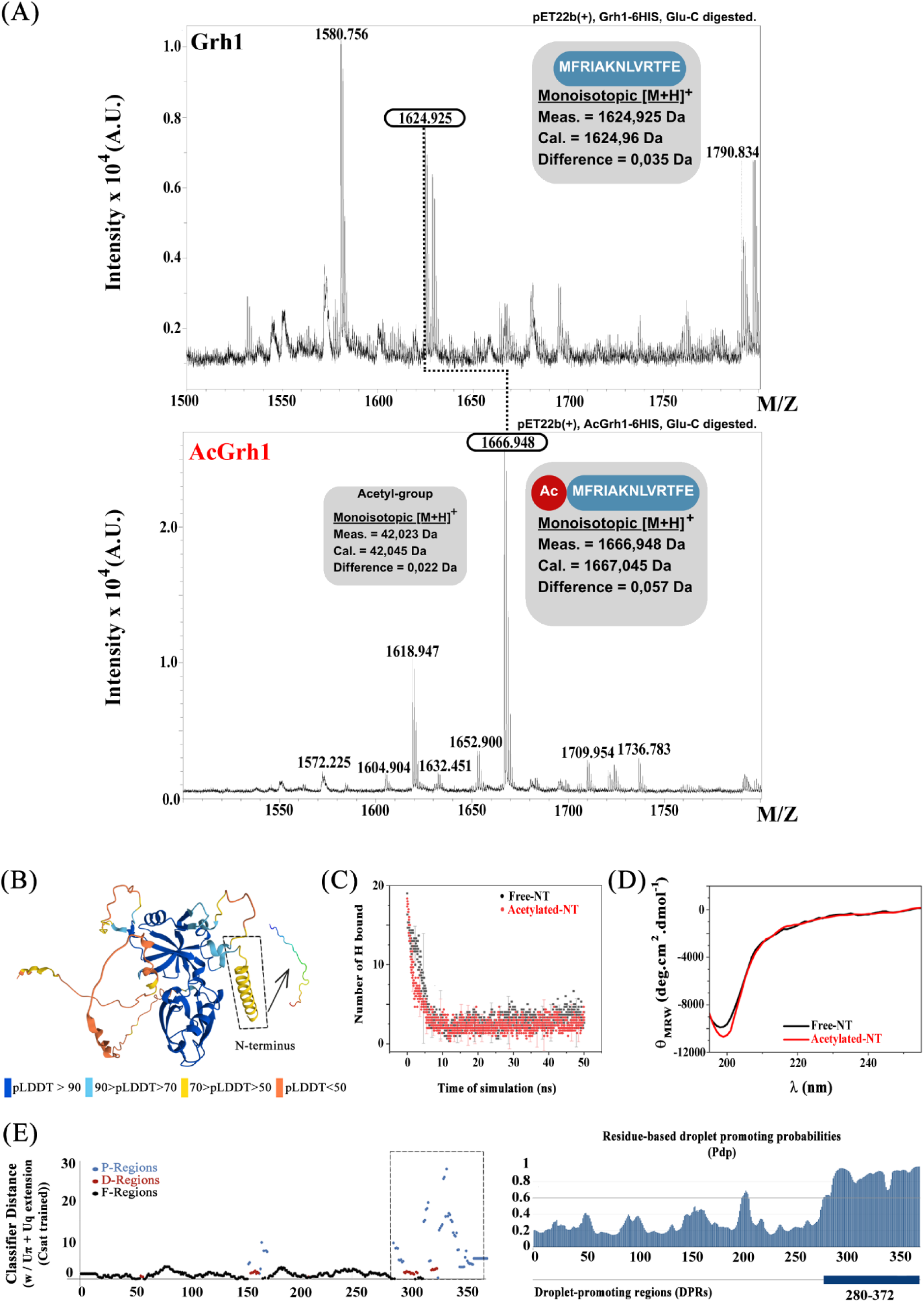
Structural features and condensation propensity of Grh1. (A) MALDI-MS analysis of GluC-digested Grh1 and AcGrh1. The spectra show the expected m/z values for the N-terminal peptides of both proteins, with the mass shift corresponding to the addition of an acetyl group in AcGrh1. Notably, no signal corresponding to the non-acetylated peptide is detected in the AcGrh1 sample, indicating near-complete N-terminal acetylation within the detection limits of the technique. (B) AlphaFold 3.0 model of full-length Grh1, coloured according to pLDDT confidence scores. The predicted N-terminal α-helix, highlighted by the dashed square, exhibits intermediate confidence and is positioned upstream of the structured GRASP domain. (C) Molecular dynamics simulation of the isolated N-terminal segment in its acetylated and non-acetylated forms. The plot shows the number of hydrogen bonds over time, demonstrating progressive destabilisation and loss of the α-helical structure predicted by AlphaFold, resulting in a disordered conformation irrespective of acetylation. (D) Far-UV circular dichroism spectra of the chemically synthesised N-terminal peptides with and without acetylation. Both spectra are characteristic of fully disordered structures, supporting the conclusion that this region does not adopt a stable secondary structure in solution. (E) Residue-level prediction of phase separation propensity using ParSe (left) and FuzDrop (right) ^46,47^. In ParSe, P-regions (P) are intrinsically disordered and prone to undergo LLPS, D-regions (D) are intrinsically disordered but unlikely to undergo LLPS, and F-regions (F) may be structured or acquire structure upon binding. In FuzDrop, droplet-promoting regions are defined as stretches of at least five consecutive residues with a droplet-promoting probability (pDP) of 0.60 or higher, highlighted as a blue bar beneath the graph. Both predictors identify the highly disordered C-terminal region as the principal driver of biomolecular condensation propensity, with no contribution from the N-terminal domain.

### Acetylation does not induce stable helical content at the Grh1 N-terminus

It is also worth noting that while Grh1 harbours a highly disordered C-terminal region, its N-terminal segment is predicted by AlphaFold 3.0 ^44,45^ to adopt a short helical conformation, consistent with previous proposals suggesting the presence of an amphipathic helix (Figure 1B). However, a short molecular dynamics simulation of acetylated and non-acetylated N-termini indicates that this helical structure is unstable in solution and rapidly unfolds, suggesting it lacks sufficient intrinsic stability (Figure 1C). To experimentally validate this observation, we synthesised the entire N-terminal segment preceding the structured GRASP domain in acetylated and non-acetylated forms and analysed their structural properties using far-UV circular dichroism (CD) spectroscopy. The resulting spectra clearly show that both constructs were fully disordered in solution, displaying CD signatures characteristic of disordered chains (Figure 1D). Furthermore, the spectra of the acetylated and non-acetylated peptides were virtually indistinguishable, providing no evidence that Nt-acetylation induced any disorder-to- order transition in this region. Despite its disordered nature in solution, multiple phase-separation predictors consistently suggest that this N-terminal region is not a significant contributor to the phase separation propensity of Grh1 (Figure 1E).

### Biomolecular condensates of Grh1 and AcGrh1

Biomolecular condensates are dynamic supramolecular assemblies formed through phase separation coupled to percolation, a process in which macromolecules spontaneously demix into coexisting dense and dilute phases ^5^. These assemblies enable spatial compartmentalisation without membranes, concentrating specific biomolecules to regulate biochemical reactions, modulate cellular organisation, and orchestrate stress responses ^1^. At the molecular level, condensation arises from a network of weak, multivalent, and transient interactions, often mediated by intrinsically disordered regions, low-complexity domains, and modular binding motifs ^4,48^. *In vitro*, phase behaviour is highly sensitive to parameters such as pH, salt concentration, temperature, and macromolecular crowding ^1^.

Previous studies from our group demonstrated that Grh1 undergoes phase separation under stress-mimicking conditions, which is potentially involved in unconventional protein secretion ^34^. However, it remained unclear whether post-translational modifications modulate the physicochemical properties of the condensates, particularly N-terminal acetylation. To address this, we investigated the condensation behaviour of recombinant Grh1 and its N-terminally acetylated counterpart (AcGrh1) under identical crowding and stress conditions. Both proteins readily formed condensates when incubated in 20 mM sodium phosphate buffer containing 10% PEG-8000 at pH 5.4, as visualised by differential interference contrast (DIC) microscopy (Figure 2A). Using the PhaseScan strategy to map the phase diagrams of both proteins across a range of PEG-8000 and protein concentrations, we observed that Grh1 and AcGrh1 respond similarly to increasing macromolecular crowding (Figure S1). As expected, higher protein concentrations promoted condensate formation in both cases, even under a fixed macromolecular crowding concentration, confirming that the saturation threshold was exceeded (Figure 2A). However, condensates formed by Grh1 were consistently more numerous than those formed by AcGrh1, indicating that N-terminal acetylation reduces nucleation frequency under otherwise identical physicochemical conditions.

**Figure 2:**
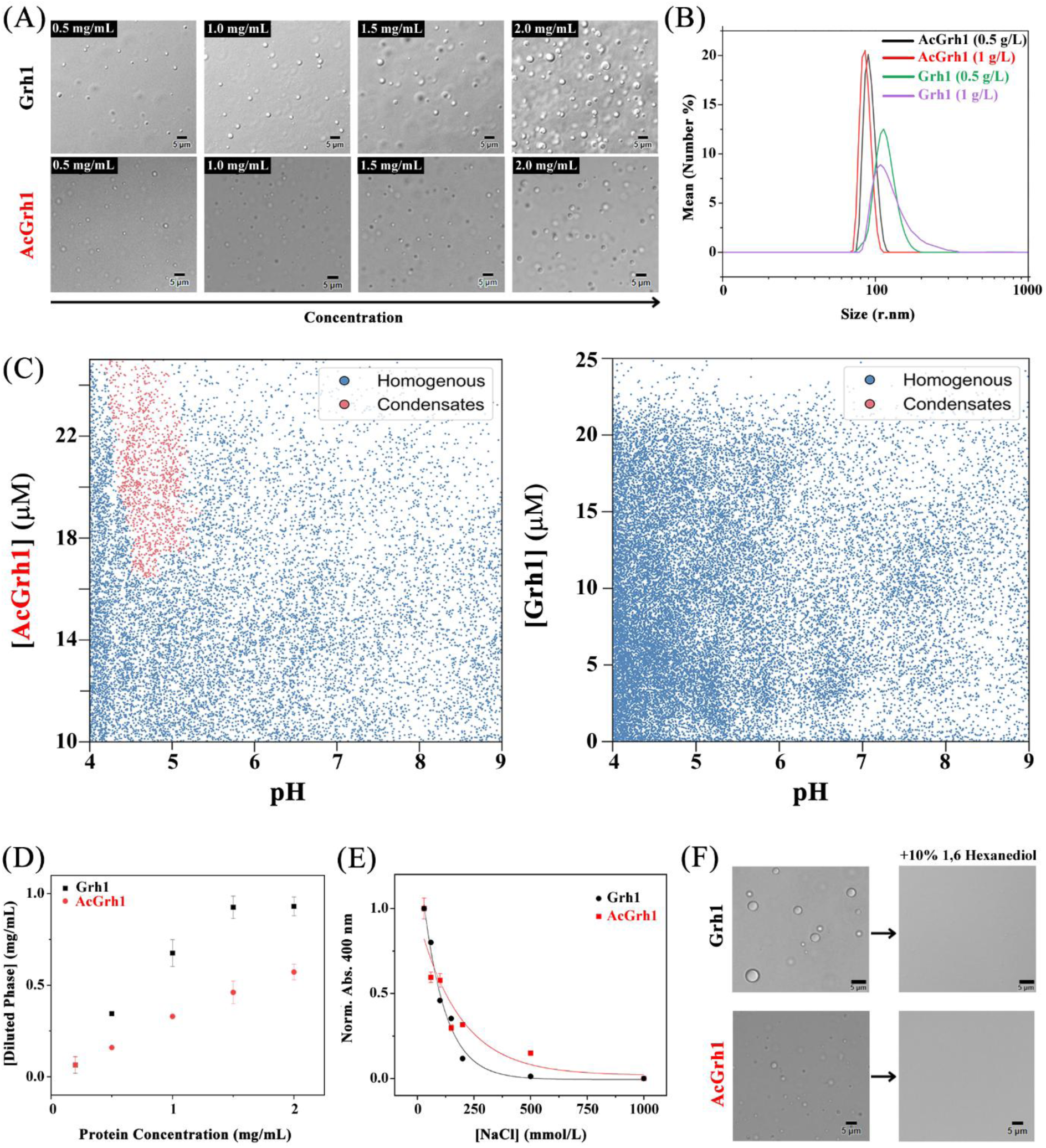
Grh1 and AcGrh1 biomolecular condensation analyses. (A) Differential interference contrast (DIC) microscopy images of condensates formed by Grh1 and AcGrh1 at increasing protein concentrations (0.5 to 2.5 mg/mL). Both proteins form condensates under crowding conditions (10% PEG 8000, pH 5.4), though Grh1 exhibits a higher number and larger condensate size than AcGrh1. (B) Dynamic light scattering (DLS) analysis of condensate size distribution at two protein concentrations (0.5 and 1 mg/mL). AcGrh1 droplets are consistently smaller and exhibit a more homogeneous size distribution than Grh1 condensates. (C) Phase diagram mapped using the PhaseScan microfluidic platform reveals protein concentration as a function of pH for Grh1 and AcGrh1, without the addition of any agent for macromolecular crowding. Each point represents a microdroplet, with blue indicating homogeneous solutions and red indicating phase-separated droplets. The acetylated form reduces the tendency to phase separate in the physiologically relevant acidic range. (D) Turbidity measurements at 400 nm of the diluted phase as a function of total protein concentration reveal a significantly lower turbidity signal for AcGrh1 compared to Grh1 across the concentration range. (E) Turbidity at 400 nm as a function of NaCl concentration. Both proteins exhibit a reduction in turbidity with increasing salt concentration, consistent with the disruption of electrostatic interactions; however, the effect is more pronounced for Grh1. (F) DIC microscopy images of Grh1 and AcGrh1 condensates after treatment with 10% (v/v) 1,6-hexanediol, a hydrophobic interaction disruptor. In both cases, the condensates are completely dissolved, indicating that hydrophobic interactions play a central role in the stability of the condensates.

Despite the shared ability to form condensates, marked differences were observed in the morphology and physical properties of the resulting assemblies. Dynamic light scattering (DLS) measurements revealed that condensates formed by AcGrh1 were significantly smaller in hydrodynamic radius compared to those formed by unmodified Grh1 (Figure 2B). This observation suggests that acetylation modifies the interaction landscape of the protein, leading to the formation of less multivalent, cross-linked assemblies, resulting in smaller, more dispersed condensates (Figures 2A and 2B).

Interestingly, several physicochemical parameters were also markedly affected by Nt-acetylation. Grh1 phase separation is tightly linked to stress responses associated with unconventional protein secretion in yeast, where cytosolic acidification has been proposed as a key regulatory signal. Given this, pH sensitivity represents a central aspect of Grh1 condensation. Although light microscopy confirmed that both Grh1 and AcGrh1 could form biomolecular condensates under various pH conditions (Figure S2), it remained unclear whether acetylation alters the pH dependence of phase separation. To address this, we employed PhaseScan, a high-throughput droplet microfluidics-based platform that enables us to systematically map phase diagrams across a pH gradient ^39^. While the experimental setup differs from microscopy-based assays, notably by eliminating the water-air interface and replacing it with an oil overlay, the resulting phase maps provide an unbiased and quantitative overview of protein condensation behaviour. The results reveal that acetylation significantly perturbs the phase diagram of Grh1, resulting in a marked reduction in condensate formation within the pH range most relevant to cytosolic stress (Figure 2C). This data suggests that N-terminal acetylation impairs the protein’s ability to respond to acidification, potentially modulating its function during stress-induced secretion.

To quantify the differences in condensation propensity between the two forms, we experimentally determined the saturating concentration (Csat) for each construct. Csat represents the critical threshold beyond which a homogeneous protein solution becomes thermodynamically unstable, triggering phase demixing into coexisting dilute and dense phases. This equilibrium point is a defining feature of phase-separated systems, reflecting the intrinsic ability of a protein to condense under specific conditions. In our experiments, solutions with different total protein concentrations were prepared at a constant ionic strength and crowding, subjected to centrifugation, and then the protein concentration in the supernatant (dilute phase) was measured. The onset of phase separation was identified as the plateau in soluble protein concentration, which indicates saturation. The results, shown in Figure 2D, reveal a pronounced difference between the two protein variants: Grh1 exhibits a substantially lower saturation point than AcGrh1. This data confirms that Nt-acetylation decreases the protein’s propensity to undergo phase separation, requiring higher concentrations to reach the full demixing threshold.

We performed turbidity assays in the presence of salt, crowding agents, and chemical disruptors to investigate further the physical nature of the intermolecular interactions that govern condensate stability. Polyethylene glycol (PEG), used here to mimic macromolecular crowding, enhanced turbidity in a concentration-dependent manner for both Grh1 and AcGrh1. However, the effect was markedly more pronounced for the non-acetylated protein, consistent with its stronger propensity to undergo phase separation (Figure S3A). Titration with NaCl led to a progressive decrease in turbidity for both protein variants, indicating that electrostatic interactions contribute significantly to condensate formation (Figure 2E). Notably, this reduction was more severe for Grh1, suggesting that AcGrh1 condensates rely less on charge-based interactions. We then introduced urea at sub-denaturing concentrations below the threshold for global unfolding to probe the role of hydrogen bonding. Urea perturbs hydrogen bonds and weakens interactions that stabilise condensed phases. Both Grh1 and AcGrh1 showed a moderate but comparable decrease in turbidity, indicating that hydrogen bonding contributes to condensate integrity in both cases (Figure S3B). Finally, we tested the sensitivity of the condensates to 1,6-hexanediol, a hydrophobic interaction disruptor commonly used to assess LLPS stability. Adding 1,6-hexanediol led to the complete dissolution of condensates for Grh1 and AcGrh1, confirming that hydrophobic interactions play a central role in stabilising the condensates in both protein variants (Figure 2F). These results indicate that N-terminal acetylation selectively modulates electrostatic contributions to condensate formation, leaving hydrogen bonding and hydrophobic interactions largely unaffected.

### Nanoscale organisation and material properties of Grh1 condensates

To further investigate how N-terminal acetylation modulates the internal organisation and material properties of condensates, we used the environmentally sensitive fluorescent probe ACDAN^49^. This dye responds to local water dynamics through solvent dipolar relaxation, providing a readout of molecular crowding within condensates ^50^. Upon excitation, ACDAN undergoes a dipole change; in water-rich environments, the emission is red-shifted due to solvent relaxation, whereas in crowded, less fluid microenvironments, the emission is blue-shifted ^50^. We performed hyperspectral imaging combined with phasor plot analysis to map the polarity and crowding profiles within condensates formed by Grh1 and AcGrh1 (Figure 3). In Grh1 condensates, we observed a clear shift in water accessibility between pH 5.4 and 7.4: acidic pH (mimicking stress) promoted higher solvent relaxation, suggesting a more hydrated and dynamic environment. In contrast, at pH 7.4, Grh1 condensates appeared more restricted and densely packed (Figure 3A). Interestingly, AcGrh1 showed similar relaxation profiles at both pH conditions, indicating reduced responsiveness to environmental pH and suggesting a more rigid, less hydrated internal organisation overall (Figure 3B).

**Figure 3:**
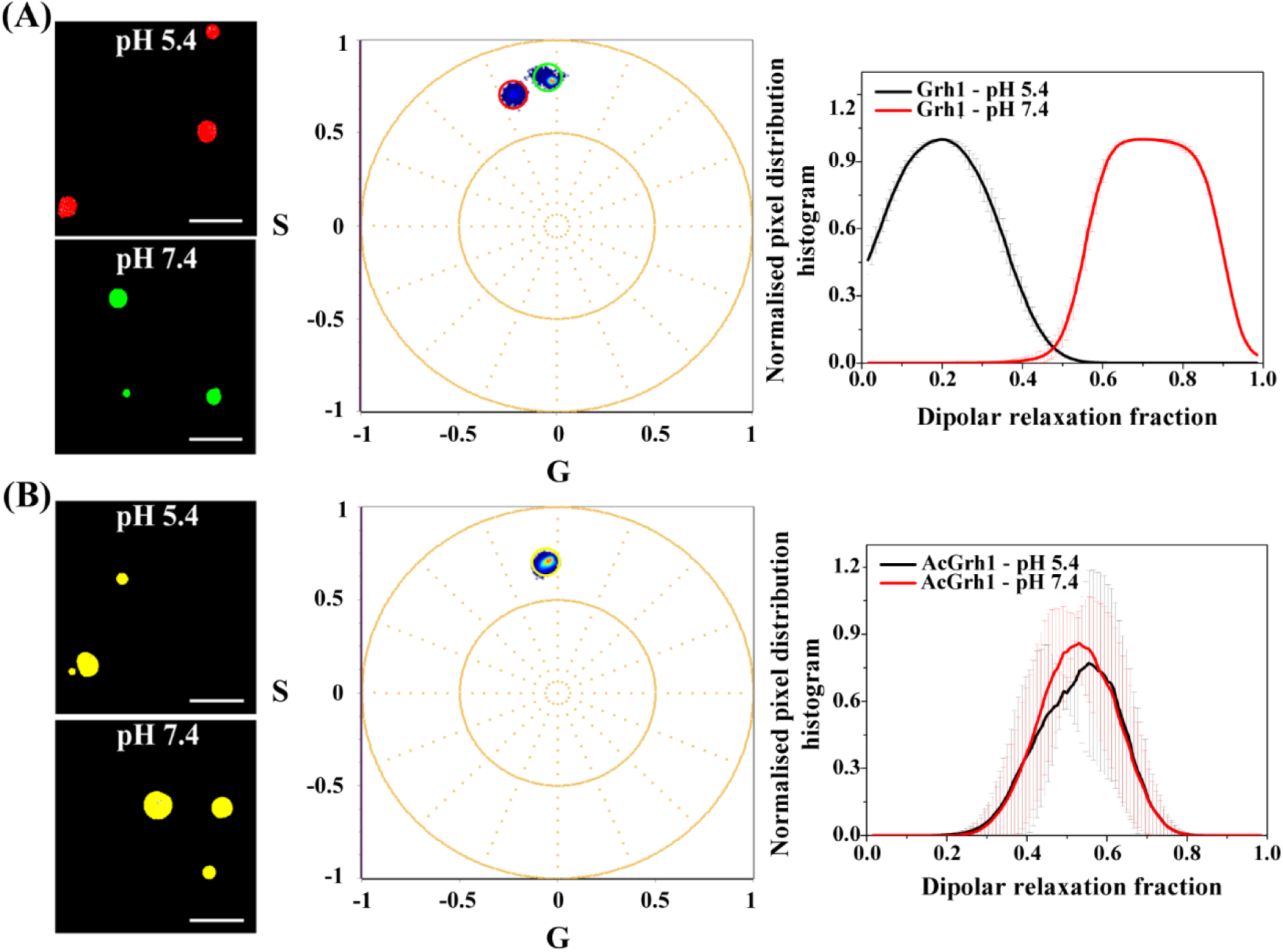
Internal hydration and environmental responsiveness of Grh1 and AcGrh1 condensates assessed by ACDAN hyperspectral imaging. Hyperspectral phasor plots of ACDAN fluorescence in condensates formed by (A) Grh1 and (B) AcGrh1 at pH 5.4 and 7.4. Each data point represents a pixel in the corresponding fluorescence image, with colour-coded cursors (yellow, red, green) indicating representative pixel populations mapped onto the spatial images. Grh1 condensates at pH 5.4 exhibit increased dipolar relaxation, suggesting higher water accessibility compared to those formed at pH 7.4. In contrast, AcGrh1 condensates display minimal spectral shifts between pH conditions, indicating reduced hydration and dampened environmental responsiveness. Dipolar relaxation fractions were calculated using a two-cursor analysis. Scale bars: 20 µm.

To further explore the material properties of Grh1 condensates and their response after Nt-acetylation, we investigated the miscibility of Grh1 and AcGrh1 within the same phase-separated assemblies. Fluorescently labelled variants of each protein were generated, and their co-condensation behaviour was assessed by fluorescence and confocal microscopy. Both proteins were readily partitioned into shared condensates at equimolar concentrations, yielding merged fluorescence signals in droplets of varying sizes (Figures 4A and B), indicating a degree of miscibility under these conditions. Closer examination, however, revealed the emergence of structured, multiphase condensates exhibiting a core-shell organisation: Grh1 localised preferentially to the condensate core, while AcGrh1 was enriched at the periphery (Figures 4A, C and D). This spatial segregation suggests that although the two forms are phase-separated within the same droplet, their differential partitioning reflects distinct interfacial properties, most likely governed by their surface tension with the surrounding medium. Such behaviour closely resembles capillarity-driven structuring observed in other multiphase condensates^10,51^. In these systems, immiscible or partially miscible protein phases can adopt stable spatial arrangements driven by the minimisation of interfacial free energy. According to thermodynamic principles, interfaces with the highest surface tension incur the largest energetic penalty and are thus minimised, favouring the internal positioning of the phase with the lowest interfacial tension ^51^. In our system, the Grh1-rich phase exhibits lower interfacial tension with the surrounding buffer, explaining its localisation at the droplet interior. In contrast, AcGrh1 potentially exhibits increased surface tension relative to the bulk solvent phase and thus accumulates at the bulk solvent/Grh1 condensate interface. Remarkably, these effects are elicited by a single chemical modification: the acetylation of the N-terminal amine. Despite the compositional similarity of Grh1 and AcGrh1, this minimal post-translational modification substantially alters the interfacial properties of the condensates, reshaping not only the morphology of individual condensates but also their internal organisation.

**Figure 4:**
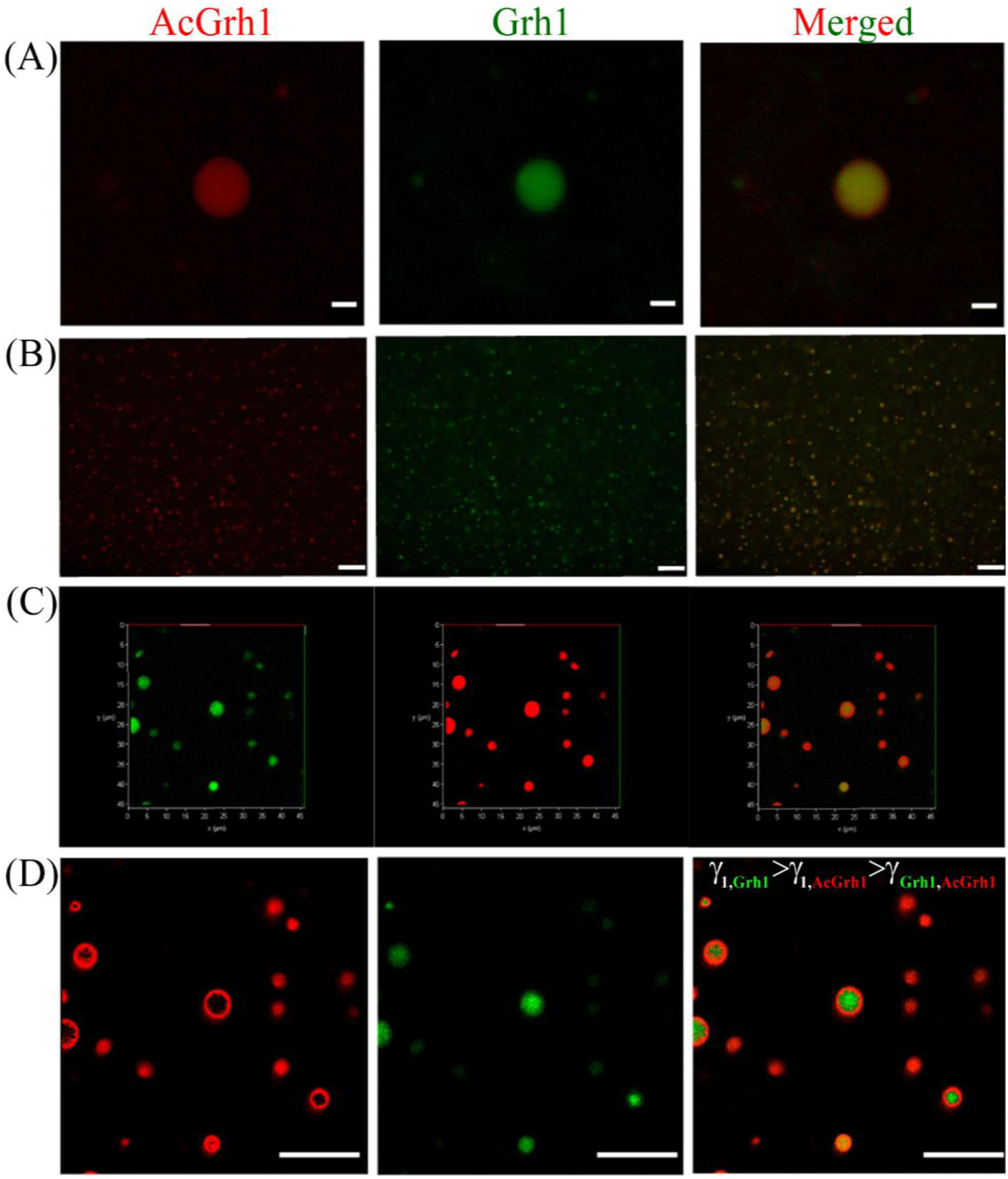
Co-condensation of Grh1 and AcGrh1 reveals the formation of multiphase condensates with core-shell organisation. (A) Fluorescence microscopy images of Grh1 (Alexa Fluor 488, green) and AcGrh1 (Alexa Fluor 546, red), both at 1 mg/mL in 20 mM sodium phosphate buffer (pH 5.4) with 10% PEG-8000. Under these conditions, both proteins form condensates, with substantial signal overlap in the merged image, indicating co-condensation. Scale bar: 5 µm. (B) Wide-field fluorescence microscopy image showing a larger population of condensates. Colocalisation remains prominent, suggesting miscibility between Grh1 and AcGrh1 across multiple droplets. Scale bar: 20 µm. (C) Confocal microscopy Z-stack cross-section of representative condensates, highlighting the internal spatial organisation. A core-shell architecture is observed, with Grh1 localised to the condensate core and AcGrh1 enriched at the periphery. Scale bar: 20 µm. (D) Confocal 3D reconstruction of representative condensates, with fluorescence intensity visualised using Look-Up Tables (LUTs) in ImageJ. This analysis confirms the spatial segregation between Grh1 and AcGrh1 and supports the presence of multiphase organisation. The observed core-shell structure reflects differences in interfacial tension: three interfacial energies are relevant—between Grh1 and the bulk phase (γ₁,_Grh1_), AcGrh1 and the bulk (γ₁,_AcGrh1_), and Grh1 and AcGrh1 (γ_AcGrh1,Grh1_). Scale bar: 10 µm.

Together, these findings demonstrate that Nt-acetylation exerts a broad and multifaceted influence on the condensation behaviour of Grh1. This single, chemically subtle modification not only alters critical parameters such as pH sensitivity and saturation concentration but also reshapes the balance of intermolecular forces, dampening electrostatic interactions, reducing effective valency, and modulating conformational dynamics. These changes impact the rheological properties of the condensates, influencing their size, density, miscibility, internal hydration, and responsiveness to environmental cues. By tuning both the nucleation threshold and the internal architecture of phase-separated assemblies, Nt-acetylation emerges as a potent regulatory mechanism capable of fine-tuning condensate behaviour without altering the underlying protein sequence.

### Exploring the impact of Nt-acetylation on Grh1 structural and biochemical features

Given the substantial impact of Nt-acetylation on Grh1 condensate formation, we next sought to explore the underlying principles that might account for these effects. We examined whether this modification altered the general biochemical and structural properties of these proteins in solution under non-phase-separated conditions. Specifically, we assessed the influence of acetylation on Grh1’s secondary structure using circular dichroism (CD) spectroscopy in the far-UV region, which is particularly sensitive to backbone conformations and serves as a reliable method for detecting disorder-to-order transitions such as α-helix formation ^52,53^. N-terminal acetylation has been shown to stabilise α-helical conformations in several proteins, including α-synuclein, where it enhances N-terminal helicity and reduces aggregation propensity ^54,55^. However, for Grh1, the far-UV CD spectra revealed no appreciable differences between the acetylated and non-acetylated forms (Figure 5A). The absence of any detectable changes in the CD spectrum indicates that acetylation does not promote helix formation under these conditions or that any structural transition is minimal and below the detection limit of this approach. This data also agrees with the MD simulations shown in Figure 1B.

**Figure 5:**
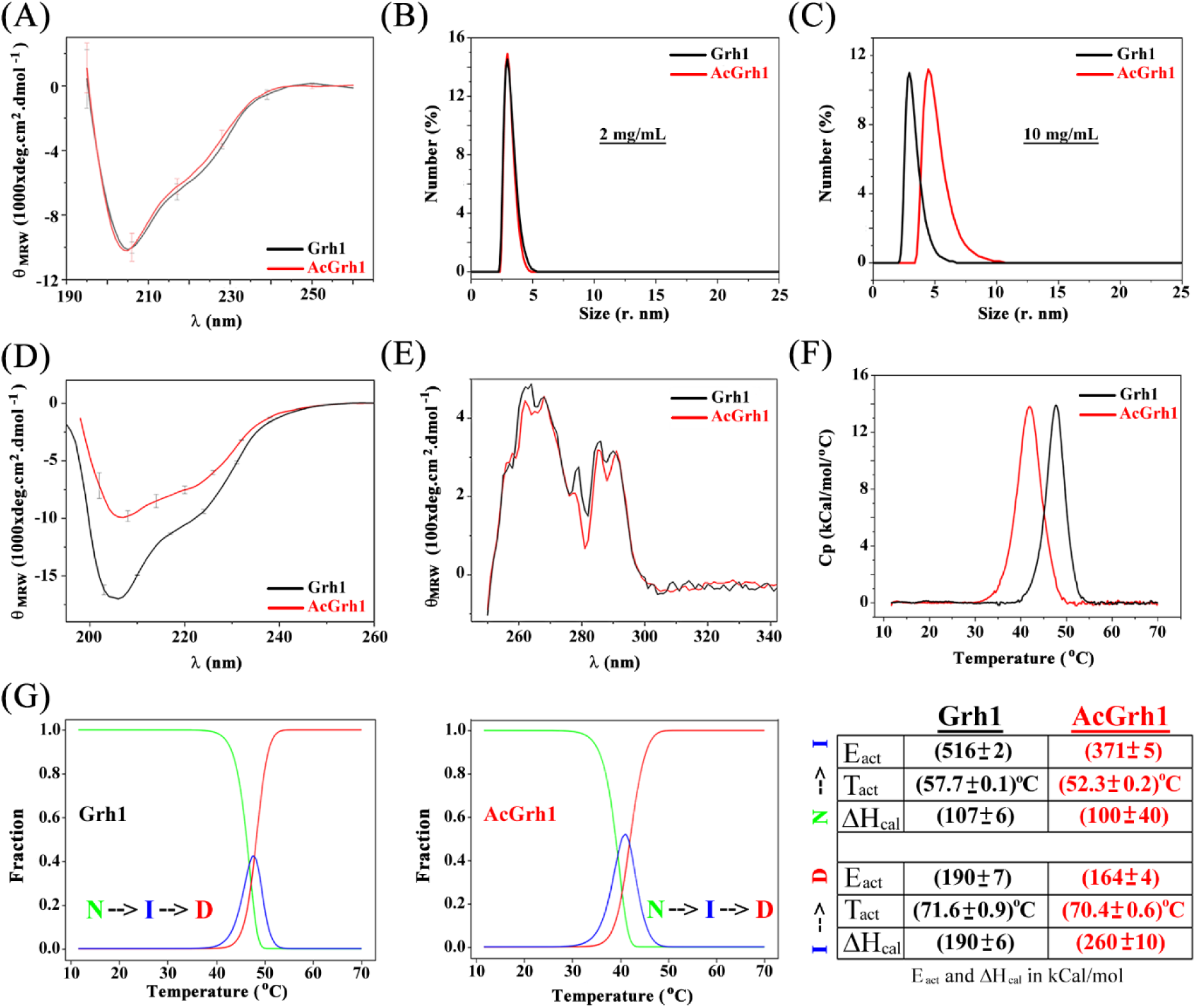
(A) Far-UV circular dichroism (CD) spectra of Grh1 and AcGrh1 at low concentration (0.1 mg/mL). Under these conditions, both proteins exhibit indistinguishable secondary structure profiles consistent with predominantly disordered or flexible conformations. (B–C) Dynamic light scattering (DLS) analyses of Grh1 and AcGrh1 at moderate (2 mg/mL) and high (10 mg/mL) protein concentrations. AcGrh1 displays a clear shift toward higher hydrodynamic radius consistent with oligomerisation at elevated concentrations, while Grh1 remains monomeric. (D) Far-UV CD spectra of Grh1 and AcGrh1 at high concentration (10 mg/mL). AcGrh1 shows a marked increase in α-helical content at this concentration, indicated by the characteristic shift of the spectral minimum from 205 to 208 nm. (E) Near-UV CD spectra probing the tertiary structure of both variants. The overall aromatic environment appears largely unchanged between the two proteins, suggesting that acetylation-induced structural rearrangements primarily occur in regions lacking aromatic residues, likely within disordered segments. (F) Differential scanning calorimetry (DSC) thermograms for Grh1 and AcGrh1 were acquired under identical experimental conditions. AcGrh1 exhibits a lower transition temperature and reduced cooperativity relative to Grh1. (G) Populations of native (N), intermediate (I), and denatured (D) states derived from fitting the DSC data to a three-state irreversible model using CalFitter. The accompanying table summarises the thermodynamic parameters for each transition step in the thermal denaturation of both proteins.

In parallel, we explored the potential effect of acetylation on protein oligomerisation. Although Grh1 is generally monomeric under physiological conditions, a functional dimeric state has been proposed in its roles at the Golgi ^35^. Moreover, the human GRASP representative GRASP55 has already been shown to form dimers only when myristoylated ^56^. To determine whether acetylation alters oligomerisation, we first employed dynamic light scattering (DLS) and size-exclusion chromatography coupled to multi-angle light scattering (SEC-MALS) at moderate protein concentrations (∼2 mg/mL), comparable to those used in the phase-separation studies shown in Figures 2, 3, and 4. Under these conditions, both Grh1 and AcGrh1 were monodisperse and monomeric, exhibiting similar hydrodynamic radii (Figure 5B and S4). These findings indicate that N-terminal acetylation does not promote self-association at intermediate concentrations. However, those concentrations are likely below the physiological range experienced by Grh1 *in vivo*. As a membrane-associated protein, Grh1 may reach much higher local concentrations at the Golgi surface, and within biomolecular condensates, protein concentrations can exceed 100 mg/mL ^1,57^. To better approximate these conditions, we performed DLS measurements at the highest concentration achievable from our purification yields (∼10 mg/mL). Remarkably, at this elevated concentration, AcGrh1, but not Grh1, displayed a clear shift toward a dimeric state, indicating that N-terminal acetylation facilitates dimerisation at high concentrations (Figure 4C). This oligomerisation event may have profound consequences for condensation. Dimerisation reduces the number of accessible interaction motifs, effectively decreasing multivalency, since residues involved in the dimer interface are no longer available for intermolecular contacts. This reduction in valency may underlie the higher saturation concentration and smaller condensate sizes observed for AcGrh1, offering a mechanistic link between acetylation-induced oligomerisation and suppressed phase separation.

Given the promiscuous role of Grh1 in unconventional secretion, Golgi organisation, and dynamics, conformational plasticity may hold functional significance. Motivated by this and the clear evidence of acetylation-dependent dimerisation, we revisited the CD measurements at high protein concentrations. By reducing the cuvette path length by two orders of magnitude (to 0.01 mm), we could perform far-UV CD spectroscopy at concentrations comparable to those used in the DLS assays. Under these conditions, a clear spectral shift was observed for AcGrh1, with the minimum at 205 nm shifting toward 208 nm, and the overall profile acquiring features typical of α-helical proteins (Figure 5D). This increase in helical content was not observed in the unmodified Grh1 (Figure 5D). Complementary measurements in the near-UV region (260–320 nm), which report on the chiral environments of aromatic residues, revealed similar spectra for both Grh1 and AcGrh1 (Figure 5E). However, subtle but reproducible differences were observed in regions corresponding to Tyr, Phe, and Trp, suggesting that tertiary structural features or side-chain packing were affected by Nt-acetylation (Figure 5E). The perturbation of secondary structure content observed in AcGrh1 at high concentrations is likely attributable to local shifts in dynamics or side-chain environments, particularly in regions depleted of aromatic residues, which are most likely flexible or disordered segments. While the precise structural elements involved remain undefined and are beyond the scope of this study, these findings provide strong evidence that Nt-acetylation induces a concentration-dependent conformational rearrangement not present in the unmodified form.

To investigate whether the observed changes in quaternary structure also affect protein stability, we next examined the thermal unfolding profiles of Grh1 and AcGrh1 using differential scanning calorimetry (DSC), performed at concentrations where dimer formation by AcGrh1 was already detectable (Figure 5F and S5). The resulting thermograms revealed a single major thermal transition for both protein variants. However, the broadened shape of the endothermic peaks suggested a multistep unfolding process. Importantly, significant differences were observed between the two proteins: AcGrh1 exhibited a ∼6 °C decrease in the main transition temperature compared to Grh1, indicative of reduced thermal stability, and the transition itself was markedly broader, consistent with decreased unfolding cooperativity. These observations suggest that Nt-acetylation increases conformational flexibility or structural heterogeneity, resulting in a less concerted unfolding.

We fitted the calorimetric data to increasingly complex models to further dissect the unfolding mechanism. As the unfolding transitions for both proteins were irreversible, a two-state irreversible model (N → D) was initially applied but failed to describe the data adequately. In contrast, a three-state irreversible model (N → I → D) provided an excellent fit in both cases (Figure 5G and S6). According to this model, Grh1 undergoes a cooperative transition through a well-defined intermediate state, sharply populated over a narrow temperature window. AcGrh1, however, populates a broader and more diffuse intermediate over a wider range, reflecting a more gradual and less ordered unfolding process. The most striking differences between the two proteins were found in the first transition (N → I). For AcGrh1, this step exhibited a ∼30% reduction in the activation energy and a 5.4 °C decrease in the activation temperature relative to Grh1, suggesting that the native state of the acetylated protein is significantly less stable and more prone to transition into intermediate conformations (Figure 5G). By contrast, only minor changes were observed for the second transition (I → D), apart from a modest reduction in the calorimetric enthalpy, implying that once the intermediate state is populated, the final denaturation step proceeds similarly in both variants. Notably, the intermediate state of AcGrh1 appears at temperatures near physiological conditions, unlike that of Grh1 (Figure 5G). This raises the possibility that the acetylated protein may sample partially unfolded or dynamic conformations under native or stress-related cellular contexts, which could be functionally relevant given its role in unconventional secretion and membrane-associated localisation.

These experiments reveal that Nt-acetylation significantly alters Grh1’s thermodynamic, structural, and oligomeric properties. Although chemically subtle, this modification reshapes the protein’s conformational energy landscape and promotes dynamic equilibria absent in the unmodified state. These findings support a broader paradigm in which post-translational modifications act as static structural regulators and fine-tuners of folding cooperativity, conformational flexibility, and condensation behaviour under physiological conditions.

## Conclusion

This study reveals that Nt-acetylation, despite modifying only a single terminal amino group, has far-reaching consequences for the phase behaviour, structure, and material properties of Grh1, a protein central to stress-induced unconventional secretion in yeast. By comparing acetylated and non-acetylated forms under controlled conditions, we demonstrate that this irreversible co-translational modification substantially reshapes multiple features of biomolecular condensation. Nt-acetylation raises the saturation concentration required for phase separation, dampens responsiveness to acidic pH, reduces droplet size and nucleation frequency, and alters the physical forces stabilising condensates, most notably by reducing electrostatic contributions. These effects are mechanistically linked to acetylation-induced dimerisation, a reduction in multivalency, and conformational changes, including an increase in α-helical content at high concentrations.

Notably, acetylation also reprograms condensate architecture and internal environment. Co-condensation experiments revealed structured multiphase assemblies with a core-shell organisation, consistent with differences in interfacial tension between Grh1 and AcGrh1. In parallel, hyperspectral imaging with the environmentally sensitive dye ACDAN uncovered striking differences in water dipolar relaxation: Grh1 condensates exhibit higher hydration and pH responsiveness, while AcGrh1 structures are more dehydrated and rigid. These results suggest that Nt-acetylation modulates not only the formation of condensates in Grh1 but also their internal solvent dynamics, an increasingly recognised determinant of condensate functionality.

Although previous studies have suggested that N-terminal acetylation has only modest effects in truncated variants of FUS ^58^, we demonstrate that this modification can profoundly impact phase separation and condensate properties in the context of a full-length protein. Together, our findings support a model in which Nt-acetylation functions as a tuneable regulator of condensate material state, integrating concentration, conformation, and interfacial energetics to control phase behaviour. Since ∼80% of human proteins are N-terminally acetylated but most recombinant studies ignore this PTM, our results underscore the need to re-evaluate experimental systems that aim to model native condensation. We propose that Nt-acetylation, often overlooked, may represent a widespread and conserved mechanism for modulating the structural and functional landscape of proteins within dynamic, membrane-free cellular compartments.

## Acknowledgements

The authors thank the Brazilian agencies *Conselho Nacional de Desenvolvimento Científico e Tecnológico* (CNPq – Grant no 306682/2018-4) and *Fundação de Amparo à Pesquisa do Estado de São Paulo* (FAPESP - Grants No. 2022/06006-0, 2023/17294-9, 2023/12516-3, 2015/16812-0 and 2023/04532-9) for the financial support and access to multiuser equipment facilities.

## Supplementary data

In the supplementary material, we have included Figs. S1-S6.

## Declaration of generative AI and AI-assisted technologies in the writing process

While preparing this work, the authors used ChatGPT-4o and Grammarly AI Writing Assistance solely to improve the language and readability of the text. After using these tools/services, the authors reviewed and edited the content as needed and take full responsibility for the publication’s content.

