## supplementary material for "The role of N-terminal acetylation on biomolecular condensation"

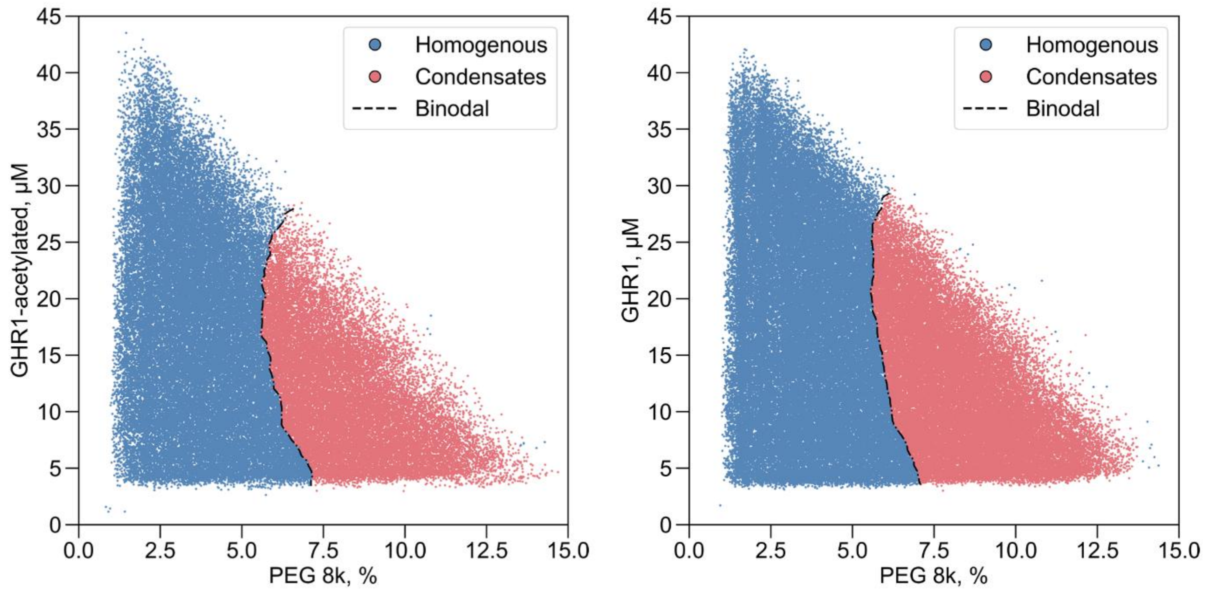


**Figure S1:** Phase diagram mapping using the PhaseScan microfluidic platform shows protein concentration as a function of PEG8000 for Grh1 and AcGrh1. Each point represents a microdroplet, with blue indicating homogeneous solutions and red indicating phase-separated droplets.

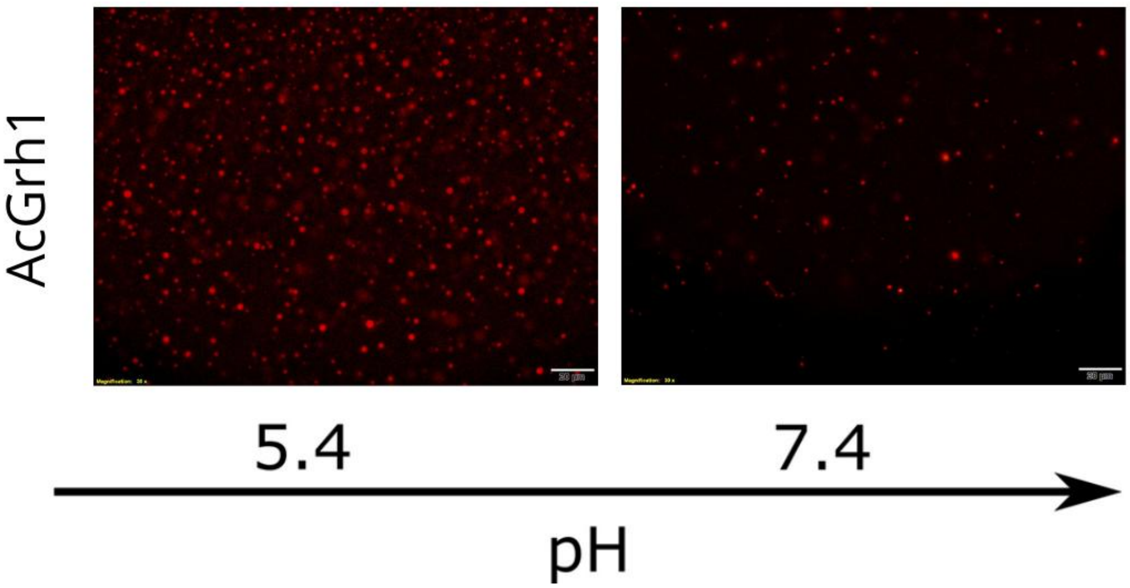


**Figure S2:** Fluorescence microscopy of AcGrh1 labelled with Alexa Fluor 546 under two physiologically relevant pH conditions. Images show AcGrh1 at pH 5.4 and pH 7.4, with 10% PEG 8000, representing cytosolic conditions associated with different functional states in Saccharomyces cerevisiae. At pH 5.4, characteristic of the acidic cytosol during starvation stress, Grh1 is involved in unconventional protein secretion. At a pH of 7.4, typical of non-stressed conditions, the protein is involved in Golgi dynamics and maintenance. Differences in condensate formation under these conditions reflect the pH sensitivity of AcGrh1 phase behaviour. Scale bar: 20 µm

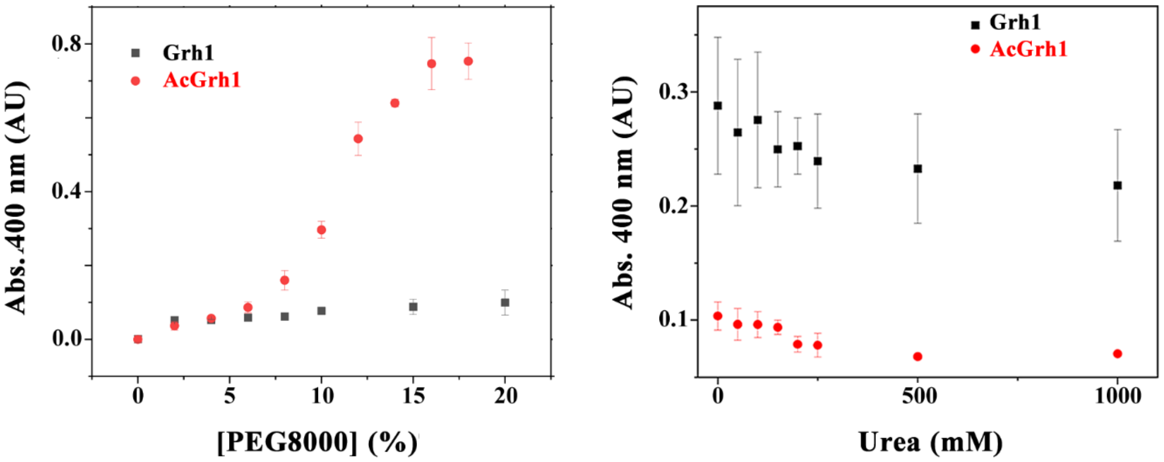


**Figure S3: Turbidity measurements at 400 nm as a function of PEG 8000 and urea concentrations for Grh1 and AcGrh1.** Absorbance at 400 nm was used to monitor condensate formation under increasing concentrations of PEG 8000 (left) and urea (right) for both Grh1 and AcGrh1. PEG was used to mimic macromolecular crowding, while urea was used at sub-denaturing concentrations to probe the role of hydrogen bonding in condensate stability.

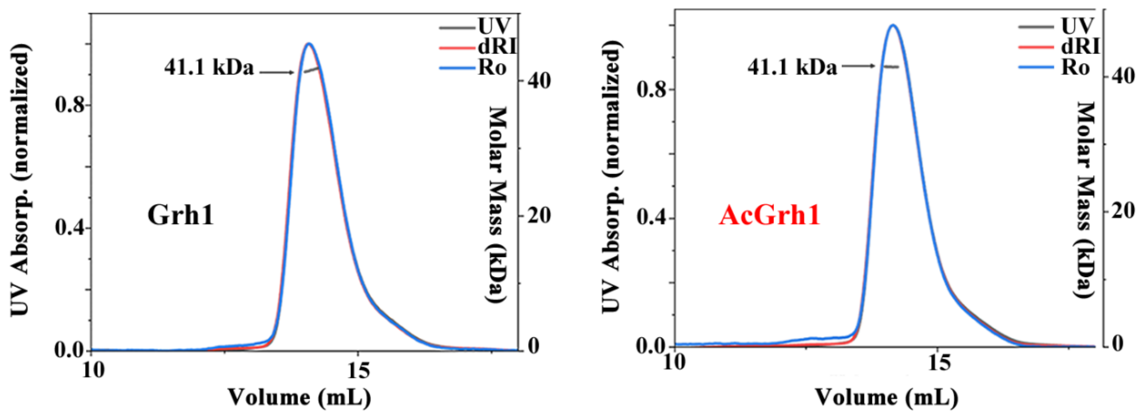


**Figure S4: SEC-MALS analysis of Grh1 and AcGrh1 at low protein concentration.** Size-exclusion chromatography coupled with multi-angle light scattering (SEC-MALS) was performed to assess the oligomeric state of Grh1 and AcGrh1 at low concentration. Both proteins eluted as monodisperse monomers, with no detectable signs of dimerisation or higher-order oligomers under these conditions.

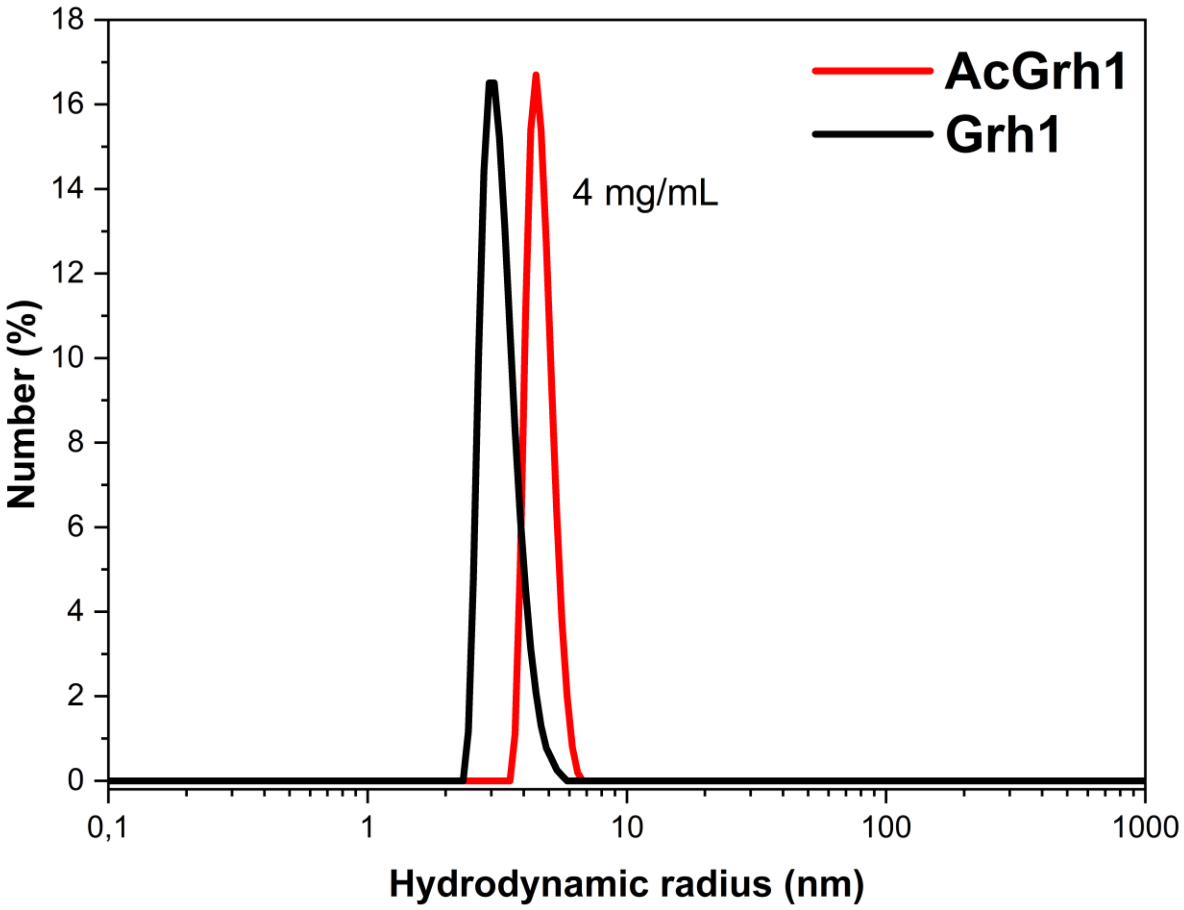


**Figure S5: Dynamic light scattering (DLS) analysis of Grh1 and AcGrh1 at 4 mg/mL.** DLS measurements were performed at 4 mg/mL to evaluate the minimal concentration at which AcGrh1 dimerisation becomes detectable, while minimising the sample consumption required for subsequent DSC experiments. At this concentration, AcGrh1 exhibits a clear shift toward a larger hydrodynamic radius, consistent with dimer formation, whereas Grh1 remains predominantly monomeric.

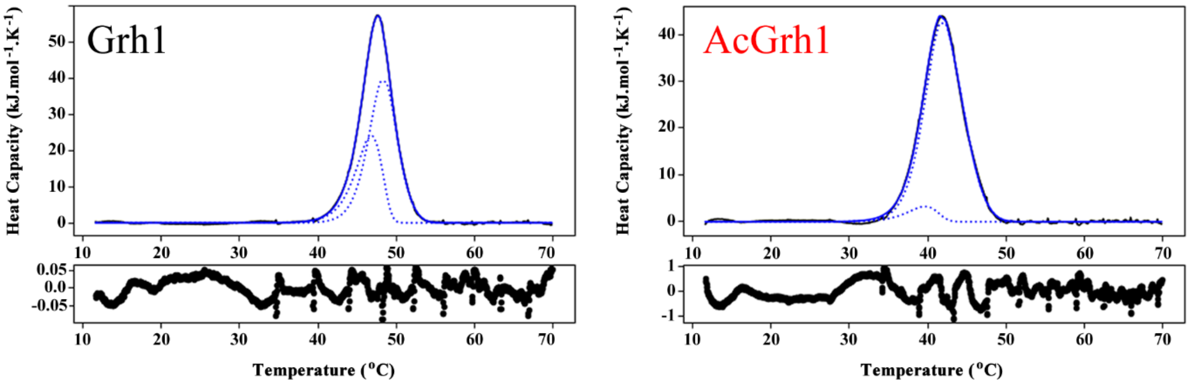


**Figure S6: Differential scanning calorimetry (DSC) fitting to an irreversible three-state unfolding model.** Experimental DSC thermograms (red) of Grh1 and AcGrh1 were fitted using an irreversible three-state transition model (N → I → D). The two individual endothermic transitions are shown as dashed blue lines, and the combined theoretical fit is shown as a solid blue line. Residuals of the fitting are displayed below each graph, indicating the quality of the model adjustment.
